# Investigating Affective and Motor Improvements with Dance in Parkinson’s Disease

**DOI:** 10.1101/665711

**Authors:** Sarah R. Ciantar, Karolina A. Bearss, Gabriella Levkov, Rachel J. Bar, Joseph F.X. DeSouza

## Abstract

**Background:** Research has supported the notion that dance alleviates motor symptoms for people with Parkinson’s disease (PD) illustrated by observed improvements in gait, balance, and quality of life. However, what remains unclear is whether engaging in weekly dance classes also positively influences nonmotor symptoms of PD, such as affect regulation (mood).

**Objectives:** To examine depressive symptoms of participants in a dance program for people with PD, and to extend previous findings on the topics for motor symptoms.

**Methods:** People with PD (n=23) and age-matched healthy controls (n=11) between the ages of 58-75 (M=67.91, SD=5.43) participated in a weekly Dance for PD^®^ class. Nonmotor symptoms of PD were assessed using the Geriatric Depression Scale (GDS), administered at three time points over the 1^st^ year of a newly-developed dance program. The Berg Balance Scale (BBS) and the Timed Up and Go (TUG) were also administered at three time points to assess motor function.

**Results:** Longitudinal mixed methods analysis showed significant improvements in GDS scores, when examining effects of the dance class over the time, with a significant main effect of time (*p* < 0.01) and condition: pre/post dance class (*p* < 0.025). Significant improvements were also observed across the motor tests of BBS (*p* < 0.001) and TUG (*p* < 0.001) measurements.

**Conclusion:** Our findings suggest dance can facilitate positive improvements in both motor and mood related symptoms of PD. These findings show important nonmotor effects of dance as an adjunct treatment for mood that may reduce the burden of this disease.

## Introduction

Parkinson’s Disease (PD) is a movement and neurodegenerative disorder estimated to affect approximately 1% of the population of adults over the age of 65 globally^1,2^. Classified as a hypokinetic disorder, PD is the product of degeneration of dopaminergic neurons in the nigrostriatal pathway of the basal ganglia, a subcortical structure associated with effortless movement (e.g. walking, speech, facial expressions). The product of dopamine dysregulation and disruption in the motor pathways in this region is impairment in the execution of movement^3^.In addition to motor impairment, various nonmotor symptoms (NMS) are also prevalent and debilitating for those with PD, such as depression, psychosis, and disordered sleep. These NMS have been shown to have a significant impact on health-related quality of life in the early stages of PD. These NMS influence factors such as institutionalization rates and health economics, which include experiences of comorbid medical conditions and health-related expense, thus playing a pivotal role in determining health and quality of life outcomes for patients with PD^4–12^.

PD is associated with a marked reduction in health-related quality of life, a measure of a patient’s subjective assessment of the impact of the disease on their physical and emotional well-being, with greater impairment seen in the advanced stages of the disease^11,13–14^. However, investigations into the influences leading to diminished quality of life in PD have shown only a weak to moderate relationship between both disease duration and severity on quality of life. Conversely, the most robust predictor, and the factor most clearly associated with worse quality of life outcomes in PD, is depression^11, 15–16^.

Depression is a frequently occurring NMS that often hallmarks the prodromal stages in PD and is associated with disease progression^17^. Past research has estimated the prevalence rate of depression to be 20-40% in people with PD (PwPD). Evidence has suggested that rather than being a product of living with a chronic condition, depression is comorbid with the pathophysiology of the disease^18, 19^. Despite the impact of depression on the overall quality of life, the affective aspect of PD is often overlooked in therapeutic approaches, which mainly target the motor symptoms of PD. In fact, the primary pharmacological intervention for PD, Sinemet (Carbidopa/Levodopa), has been suggested to exacerbate depressive symptoms^20–23^.

Due to the limitations of traditional medication-based interventions, including side effects and decreased efficacy over-time, additional adjunct treatments may be beneficial for PD patients when used in combination with medication. Various forms of augmented treatment for PD, including physical therapy, occupational therapy, exercise, tai-chi, dance, yoga, massage, acupuncture, mindfulness, meditation, herbal medicine, holistic dietary changes, and cannabis, have been gaining support in recent literature^24–32^. Findings from these supplementary interventions have primarily focused on the motor symptoms of PD, however, what remains unclear is whether an augmented treatment, such as engaging in dance, can also assist in alleviating the NMS of PD, such as depression.

Dance is a multisensory form of training and one such augmented treatment in PD which is presenting promising findings. While the mechanism of the benefits of dance in PwPD is unknown, initial research has shown neuroplastic changes associated with learning novel choreography in sub-cortical regions associated with PD (basal ganglion circuit)^30^. Preliminary research has suggested dance is an effective supplemental intervention which can improve both motor and NMS of PD^31^. Investigations of dance in PwPD have shown improvements in their performance on motor tests and mood assessments associated with regular attendance in dance classes over intervals ranging from 12-16 weeks, when compared to baseline measurements^24, 31^.

Additional feasibility research examining the effects of 4 weeks of dance on mood-specific measurements among participants with PD found a significant reduction in reported symptoms^32^. While these studies show promising results, they are limited by small sample sizes and a short duration of data collection. Additionally, how long these effects are sustained has yet to be researched, particularly for NMS such as depression. Furthermore, changes in depression in PwPD, associated with regular participation in a PD-dance intervention has yet to be examined with a measure specific to this senior population and depression in PD, such as the Geriatric Depression Scale (GDS)^33–36^.

In the present study, we examined the effect of attending weekly dance classes on NMS severity, specifically depression, in PwPD. Additionally, this study aimed to expand upon past findings regarding dance’s impact on the motor symptoms in PwPD by examining both longitudinal (1-year) and single class (1.25-hr) effects. We hypothesize that engaging in dance as a form of motor learning can act as a buffer for the neurodegeneration seen in PwPD and this will be visible through behavioral measures and a reduction in observed motor and NMS.

## Materials and Methods

### Participants

34 participants (N=34) were recruited from a Dance for PD^®^ (DfPD) program at Canada’s National Ballet School (NBS). Of these participants, 23 were diagnosed with PD between the ages of 52-76 (*M*_age_ = 67.78, *SD* = 6.14; *n*_*Males*_= 9; *M*_*DxDuration*_= 5.56 years, range = 0–17 years) and 11 were healthy controls (HC) between the ages of 61-83 (*M*_*age*_= 70.11; *SD* = 7.4; *n*_*Males*_= 6). The PD participants in the sample presented disease progression ranging from asymptomatic to severe, as measured by the H&Y scale (0–4; *M*_H&Y_ = 0.8). Written informed consent was obtained using an approved protocol from York University’s Ethics Board (2013-211).

### Measures

The GDS was used to investigate changes in the NMS of PD^33, 34^. GDS is a preferred scale used to assess depression among older populations, due to its simple format – having a dichotomous “yes/no” answering design – and its strategic omission of the somatic symptoms of depression (e.g. fatigue and psychomotor retardation) which are commonly seen in elderly people and PwPD who are not depressed^33–36^. Participants completed the GDS before and after a single dance class (1.25 hrs) at three separate time points over the six months (March, April, June; see Table 1). Data was collected at early, middle, and late phases of the dance term to capture change over time.

Motor symptoms were examined at three time points, using the Berg Balance Scale (BBS) and the Timed Up and Go (TUG) measures. The BBS is an assessment of balance and postural abilities with strong internal consistency and inter/intra-rater reliability, which is validated in PD populations^37, 38^.

The TUG measure is a subcomponent of BBS, commonly used in the clinical assessment of balance and mobility in older adults, with measurements used to infer risk of falling^37^. TUG is validated for the assessment of symptoms in PD and has been shown to be sensitive to changes in PD^40, 41^. TUG and BBS were also administered at three time points over the six months (January, April, June). Measures were collected at various time points throughout the dance term (see Table 1 for a breakdown of when each measure was collected).

### Procedures

Participants engaged in weekly 75-minute dance classes over a span of six months, during which data was collected for each of the measures three times, at four time points (January, March, April, June; see Table 1). Dance classes followed the structure of DfPD^®^, a specialized program for PwPD, which is used internationally in this demographic. Classes would begin with a seated component, followed by mirroring and paired exercises, and ending with a standing portion during which participants execute sequences of dance techniques and move across the room (see Table 2^24^). In this model, dance instructors are trained to ensure that the class is appropriate and beneficial for PwPD. Participants learned novel choreography, which was presented publicly on three separate occasions: Sharing Dance day, City Hall, and at Parkinson’s Central meetings.

Participants completed BBS and TUG before and after (pre/post) a dance class at the start of the dance term (the first time point). These tests were repeated at the third and fourth time points. The motor measurements were recorded at each time point before and after the dance class. Adhering to this data collection procedure allowed for both short-term (pre/post) and long term (January-June) comparisons. The GDS was also administered before and after the 75-minute dance classes at three points spanning the six months. Following the dance term, follow up measures for the GDS were collected after 3-months of inactivity (September) and 3-months after restarting the weekly dance classes (December), however these were stand-alone measurements that don’t follow the pre/post design.

## Results

We analyzed the data employing a linear mixed effects model analysis to account for individual subject data variability and allow for inclusion of data from participants with differing or missing data points across time. This data was analyzed in SPSS statistical software (version 24.0)^42–44^.

### Geriatric Depression Scale – GDS

Statistical analyses showed improvements in depression/mood scores when examining for the effects of the dance class over time. There was a significant main effect for time [*F* (4, 135.319) = 3.677, *p* < 0.01] and the measured effect of the dance class on GDS (pre/post) [*F* (1, 125.974) = 5.266, *p* < 0.025]; no significant interaction was found between experience (time in months dancing) and the GDS impact following a 75-minute dance class [*F* (2, 125.857) = 0.166, *p* = 0.848].

Figure 1a shows the significant reduction of GDS scores across time (March and April) and before and after the dance class for three-months of data collection. Further examination of the pre/post class comparisons used paired *t*-tests to examine the significance of the conditions (pre/post) across the differing time points and found a significant reduction in reported GDS symptoms. Significant differences were observed among the PwPD group; when examining GDS scores for the condition of the class at each of the time points as well looking at the condition between specific time points (see Supplemental Table 1a for further details on analysis).

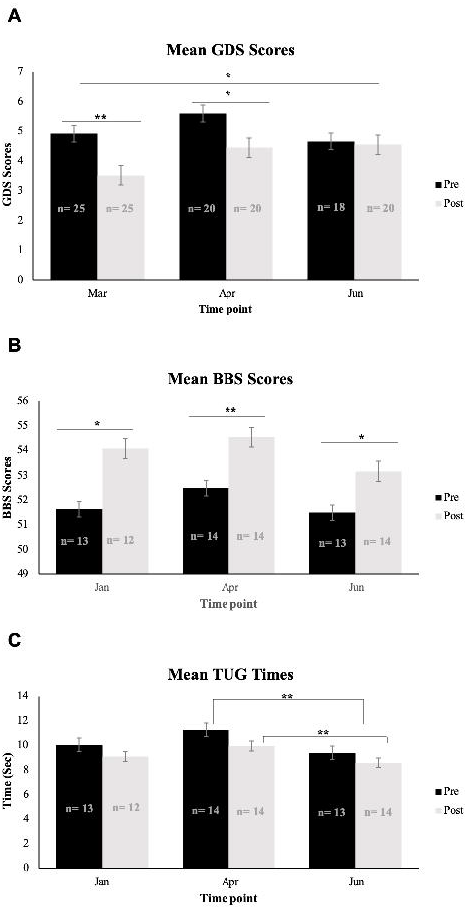

When examining the follow up measurements, a significant increase in GDS scores was observed when compared to during the dance term (See Figure 2a and 2b). This significant increase resulted in an increase in trajectories for GDS, which were previously negative during the dance term, as shown in Figure 2b.

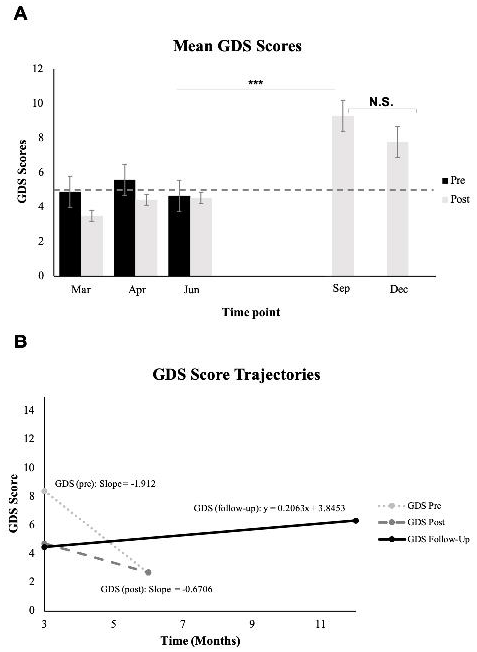

### Berg Balance Scale – BBS

BBS analysis showed significant differences for the pre/post condition [*F* (1, 60.053) = 31.346, *p* < 0.001], but not for the measurement time point [*F* (2, 64.194) =0.933, *p* = 0.398]. Examinations of an interaction between time point and pre/post condition were also not significant [*F* (2,60.051) = 0.987, *p* = 0.379]. BBS results specific to the PwPD group showed significant differences in pre/post class comparisons of scores [*F* (1, 47.397) = 29.078, *p* < 0.001]. These significant differences were also seen among the HC group for pre/post condition [*F* (1, 12.453) = 4.866, *p*< 0.05] (see Figure 1b).

### Time-Up and Go – TUG

TUG results obtained from all participants showed significant differences across the three measurement time points [*F* (2, 64.042) = 14.158, *p* < 0.001], as well as for pre/post condition comparisons [*F* (1, 61.027) = 5.317, *p* <0.05]. No significant interaction was observed between the measurement time point and the condition [*F* (2, 61.024) = 0.324, *p* = 0.724].

Further examination of TUG showed significant differences in both participant groups. When analyzing TUG times for just the PwPD group, there were significant differences between the measurement time points [*F* (2, 49.893) = 9.516, *p* < 0.001] but not across the condition of the dance class [*F* (1, 48.119) = 1.304, *p* = 0.267]. For the HC group, a significant reduction in time was seen for both the measurement time point [*F* (2, 15.232) = 15.282, *p* < 0.001] and the pre/post class condition [*F* (1, 12.585) = 21.049, *p* = 0.001]. Additionally, analyses were conducted to investigate the differences in TUG times among the measurement times. Using a pair-samples *t*-test found significant differences when comparing performance from April to June (Figure 1c; for a further breakdown of these results, see Supplemental Table 1c).

## Discussion

Our study provides novel insights into improvements in the NMS of depression, seen both short-term and long-term in PwPD, and suggest that this effect stems from engagement in weekly dance classes. To our knowledge, this is the first study revealing the benefits of dance for NMS of PD using GDS longitudinally, specifically depression^45, 46^. The lack of statistical interaction between time and condition speaks to the plausibility of there being different underlying neural processes for the immediate (condition) and long-term (time) effects of the dance class on motor and NMS. Additionally, the different effects of the dance class seen for motor as compared to NMS suggest different underlying neural mechanisms. This is also in line with neurostimulation research findings which showed a significant benefit in motor, but not mood symptoms, in PwPD with depression^47^

The results showed that for PwPD, their depression was reduced while attending the weekly dance classes, both at the level of the individual dance class as well as over time. Notice in Figure 1a that the difference between the scores in each month gets smaller over the three months, with the last sample in June showing no difference before and after the dance class, highlighting a reduction in reported GDS scores.

One noteworthy finding in the analysis of GDS scores over the term is the lack of significant interaction between the condition (pre/post) and the measurement time point. This result suggests that these benefits are independent of one another and represent two pathways for different symptom improvement, seen at both the immediate (single class) and long-term levels of dancing. These findings are important; while depression is a common occurrence in PwPD and has been shown to be predictive of disease onset, adequate interventions that target this element of the disorder are under-investigated^48^.

Interestingly, the largest support for dance as an adjunct treatment for the NMS of depression comes from the data collected at the follow-up time points – three and six months after the June measurement, and after a long summer break for Canadian summer holidays (June to September for 12weeks). GDS scores dramatically increase during the three-month period over which regular weekly dance classes were not offered, as shown in Figure 2a and 2b.

Past research examining dance movement therapy for clinical depression failed to find significant effects on the reduction of depressive symptoms^49^. This suggests that the effects on mood regulation experienced by our sample could reflect the potential benefits of dance for depression specific to PwPD.

Benefits were also observed in the motor symptoms of PD. In line with similar research on the topic, BBS scores increased, suggesting higher balance and motor abilities. Times for performing the TUG task were also reduced after a single dance class. Additionally, TUG performance improved when comparing performance taken between midpoint of the dance term (time point 3) to the last measurement time point. These findings also serve to extend and further support existing literature on the benefits of dance on the motor symptoms of PD^24–25; 45–46^. These findings are consistent with what has been shown in past research examining PwPD who participate in dance classes who showed significant improvement on TUG and BBS measures when compared to various control groups^31^.

Past research investigating dance class effects on TUG performance has produced mixed results, with some studies failing to show significant improvements^50^. While these benefits were observed at the single-class level, and in some instances when comparing between the time point measurements, there was not a significant main effect of time in BBS overall. This suggests that improvements in BBS measures were specific to the immediate effects of the individual dance class, rather than a gradual effect of the dance class over time. While this contrasts previous research, which has shown longitudinal benefits of dance for PD, the lack of significant findings may suggest neuroprotective aspects of engaging in the dance class when taking the degenerative nature of the disease into consideration. The fact that there was not a significant decrease in scores could suggest that dance is providing a protective factor to the disease^3^.

### Conclusions

Engaging in weekly dance classes over 1 year has the potential to benefit motor and NMS of PD, with visible improvements in mood regulation, balance, and postural stability. These findings are limited due to a non-active comparison group. Future research should aim to address this. Additionally, given the novel findings of the increase in depression scores following an absence of weekly dance classes, further investigations should take measurements during a break in regular dance routine to capture how long mood regulation benefits last.

Given that our findings suggest distinct pathways of improvement for motor and NMS in PD, future research should investigate the neural mechanisms involved in these changes. Through this examination, insights can be made pertaining to both the neural pathogenesis of the disease and the biological changes that neuro-rehabilitation therapies, such as dance, promote, which leads to symptom reduction.

Findings from the present study provides further evidence on the benefits of dance in PwPD for symptom reduction when engaging in weekly dance classes. A significant detriment of the burden associated with PD is the reduced quality of life, which is potentially produced by participants’ being depressed, lethargic, or unable to leave home. Through engaging in weekly dance classes, PwPD can maintain an active and social lifestyle which might further contribute to improvements in quality of life for participants. Dance therapy might further consolidate participants’ reduction in depressive symptoms and combat the stigma associated with the disease by communal class engagement. With this evidence, better-informed clinical interventions and recommendations can be tailored towards the symptoms (motor or NMS) that clinicians are seeking to alleviate.

## Author’s Roles

Contribution towards the conception and design of the study (JFXD; GL, RB), interpretation of data (GL, SC, JFXD), preparation, drafting, and revision of the manuscript (SC, JFXD, KB), and finally all authors approve of the current version for submission.

## Appendix A: Table 1

**Table 1.** Breakdown of time points for collection of measures.

| Time (Month) | Measures collected |
| --- | --- |
| 1 (January) | BBS, TUG (Pre/Post) |
| 2 (March) | GDS (Pre/Post) |
| 3 (April) | BBS, TUG, GDS (Pre/Post) |
| 4 (June) | BBS, TUG, GDS (Pre/Post) |
| 5 (September) | GDS (Follow-up) |
| 6 (December) | GDS (Follow-up) |

## Appendix B: Table 2

**Table 2.** Breakdown of a typical dance class at NBS adapted from Bearss et al., 2017.

| **<u>EXERCISE</u>** | **<u>DESCRIPTION</u>** | **<u>PURPOSE</u>** |
| --- | --- | --- |
| Dance name introduction | Stating your name with a corresponding dance movement. The rest of the class first watches before repeating the participants name and movement. Standing or seated. | Feeling welcomed and welcoming everyone in the class. Practicing skills of choreographing on the spot. |
| Tendus | Pressing the feet along the floor until the leg is fully extended. Arms follow a similar extension motion. Seated. | Warming up the feet and lower leg, while working on strengthening the core. |
| Shuffle dance | A series of shuffles, stamps, and ankle inversions. Seated. | Facilitating flexibility and mobility in the ankles and knees. |
| Magic dance | Dancing with an imaginary ball and scarf, while exploring a range of motion. Seated. | An opportunity for vivid imagery and creative interpretation. |
| Rainfall cannon | Simulating the sounds of an approaching rainstorm using various body parts as percussion instruments. Seated. | Practicing movement initiation by waiting to execute a movement in proper sequence. |
| Winning the poker game | Rising slowly from a chair while moving in a celebratory manner. | Practicing rising from a seated position in a safe manner. |
| Painter and Sculptor mirrored pairs | A paired improvisation dance, done face to face. One partner would lead while the other mirrored their painting motion. This dance finished with a series of intertwined sculpture-like poses. Seated and standing aspects. | Mirroring a partner in a detailed fashion and practicing creative movement initiation by improvising and developing unique poses. |
| Plié's in parallel and second position | Holding on to the back of a chair, plie's (bending of the knees) and rises were done in parallel (feet together) and apart. Standing. | Developing strength and balance while standing and increasing range of motion in the legs. |
| Lunging side to side | While holding onto the back of the chair, transferring weight from side to side with legs in a wide pronated position and "brandishing a fist" at a neighbouring participant. Standing. | Finding a core centre for balance by lunging off balance and returning to a central position. |
| Waltz | Waltz step performed first on the spot and then travelling. Standing. | Safely dancing through space, and physically embodying the triplet rhythm of a waltz. |
| Shy to confident shuffle dance | A standing variation of the seated shuffle dance, where the movements are done first in a demur and small manner, but gradually increase in confidence until they are gregariously expressed. | A fun way of practicing moving with confidence and with clear intention. |
| The "Showdown Hoedown" dance | Approximately a 2-min choreography done facing a partner, first dancing as adversaries in the "showdown" and then together as companions in the "hoedown." Standing. | Challenging participants to recall a lengthy piece of choreography with multiple sections and changes of direction |

