## Supplemental Table 1 for "Investigating Affective and Motor Improvements with Dance in Parkinson’s Disease"

**Appendix B: Supplemental Tables**

Supplemental Table 1a. *Pre/Post Dance Class GDS Scores.* Denote significance at the	0.05 (*) and 0.01 (**) level.

| Time | Group | *t* | *p* | *df* | Mean (*SD*) |
| --- | --- | --- | --- | --- | --- |
| Time 1: Pre/Post | PD | 3.154 | **0.007**** | 14 | 2.4 (2.947) |
|  | HC | -1.732 | 0.182 | 3 | -0.5 (0.577) |
| Time 1 Pre – Time 2 Pre | PD | -1.707 | 0.119 | 10 | -2.182 (4.238) |
|  | HC | 0.378 | 0.742 | 2 | 0.333(1.528) |
| Time 1 Pre – Time 2 Post | PD | -1.131 | 0.301 | 6 | 0.308 (4.768) |
|  | HD | 1.732 | 0.225 | 2 | 1.0 (1.0) |
| Time 1 Post –Time 2 Pre | PD | -2.643 | **0.023*** | 11 | -3.83 (5.02) |
|  | HC | -0.389 | 0.717 | 4 | -0.4 (2.302) |
| Time 1 Post – Time 2 Post | PD | -1.168 | 0.262 | 11 | -1.4 (4.641) |
|  | HC | 2.138 | 0.099 | 4 | 0.8 (0.837) |
| Time 1 Pre –Time 3 Pre | PD | -1.131 | 0.301 | 6 | -1.143 (2.673) |
|  | HC | -- | -- | -- | -- |
| Time 1 Pre –Time 3 Post | PD | 0.594 | 0.564 | 11 | 0.417 (2.429) |
|  | HC | -- | -- | -- | -- |
| Time 1 Post – Time 3 Pre | PD | -1.809 | 0.113 | 7 | -3.0 (4.69) |
|  | HC | 5.00 | 0.126 | 1 | 2.5 (0.707) |
| Time 1 Post – Time 3 Post | PD | -1.866 | 0.087 | 12 | -1.769 (3.419) |
|  | HC | -- | -- | -- | -- |
| Time 2: Pre/Post | PD | 2.679 | **0.021*** | 11 | 2.833 (3.664) |
|  | HC | 1.500 | 0.208 | 4 | 1.20 (1.789) |
| Time 2 Pre – Time 3 Pre | PD | 1.177 | 0.292 | 5 | 1.833 (3.817) |
|  | HC | 2.000 | 0.295 | 1 | 4.00 (2.828) |
| Time 2 Pre—Time 3 Post | PD | 2.482 | **0.035*** | 9 | 2.9 (3.695) |
|  | HC | 2.333 | 0.258 | 1 | 3.5 (2.121) |
| Time 2 Post – Time 3 Pre | PD | -0.950 | 0.379 | 6 | -2.0 (5.568) |
|  | HC | -- | -- | -- | -- |
| Time 2 Post – Time 3 Post | PD | -0.064 | 0.950 | 10 | -0.091 (4.70) |
|  | HC | 1.000 | 0.500 | 1 | 0.500 (0.707) |
| Time 3: Pre/Post | PD | 1.635 | 0.133 | 10 | 1.364 (2.767) |
|  | HC | 0.333 | 0.795 | 1 | 0.500 (2.121) |

Supplemental Table 1b. *Pre/Post Dance Class BBS Scores.* Denote significance at the 0.05 (*) and 0.01 (**) level.

| Time | Group | *t* | *p* | *df* | Mean (*SD*) |
| --- | --- | --- | --- | --- | --- |
| Time 0: Pre/Post | PD | -2.327 | 0.053 | 7 | -1.81 (2.20) |
|  | HC | -1.000 | 0.500 | 1 | -2.50 (3.53) |
| Time 0 Pre – Time 1 Pre | PD | 1.008 | 0.360 | 5 | 0.67 (1.63) |
|  | HC | 1.000 | 0.500 | 1 | 0.50 (0.71) |
| Time 0 Pre – Time 1 Post | PD | -1.131 | 0.571 | 5 | 0.31 (4.77) |
|  | HD | -1.000 | 0.423 | 2 | -1.67 (2.89) |
| Time 0 Post –Time 1 Pre | PD | 3.334 | **0.016*** | 6 | 2.86 (2.27) |
|  | HC | -- | -- | -- | -- |
| Time 0 Post – Time 1 Post | PD | 1.220 | 0.268 | 6 | 1.14 (2.48) |
|  | HC | -- | -- | -- | -- |
| Time 0 Pre –Time 2 Pre | PD | 1.857 | 0.137 | 4 | 1.80 (2.17) |
|  | HC | -- | -- | -- | -- |
| Time 0 Pre –Time 2 Post | PD | -0.523 | 0.629 | 4 | -0.80 (3.42) |
|  | HC | -- | -- | -- | -- |
| Time 0 Post – Time 2 Pre | PD | 3.780 | **0.013*** | 5 | 3.33 (2.16) |
|  | HC | -- | -- | -- | -- |
| Time 0 Post – Time 2 Post | PD | 1.112 | 0.317 | 5 | 0.83 (1.83) |
|  | HC | -- | -- | -- | -- |
| Time 1: Pre/Post | PD | 2.679 | **0.005**** | 11 | 2.83 (3.66) |
|  | HC | -1.000 | 0.500 | 1 | -0.50 (0.71) |
| Time 1 Pre – Time 2 Pre | PD | 1.306 | 0.228 | 8 | 0.99 (2.29) |
|  | HC | -- | -- | -- | -- |
| Time 1 Pre—Time 2 Post | PD | -0.862 | 0.414 | 8 | -0.89 (3.10) |
|  | HC | -- | -- | -- | -- |
| Time 1 Post – Time 2 Pre | PD | 3.748 | **0.006**** | 8 | 2.78 (2.22) |
|  | HC | -- | -- | -- | -- |
| Time 1 Post – Time 2 Post | PD | -0.064 | 0.606 | 9 | -1.20 (7.09) |
|  | HC | -- | -- | -- | -- |
| Time 2: Pre/Post | PD | -2.765 | **0.020*** | 10 | -2.09 (2.51) |
|  | HC | -- | -- | -- | -- |

Supplemental Table 1c. *Pre/Post Dance Class TUG Scores.* Denote significance at the 0.05 (*) and 0.01 (**) level.

| Time | Group | *t* | *p* | *df* | Mean (*SD*) |
| --- | --- | --- | --- | --- | --- |
| Time 1: Pre/Post | PD | -0.109 | 0.916 | 7 | -0.04 (0.91) |
|  | HC | 2.087 | 0.284 | 1 | 1.44(0.98) |
| Time 1 Pre – Time 3 Pre | PD | -1.930 | 0.111 | 5 | -0.95 (1.21) |
|  | HC | -3.270 | 0.189 | 1 | -2.67(1.15) |
| Time 1 Pre – Time 3 Post | PD | -1.827 | 0.127 | 5 | -0.87 (1.17) |
|  | HD | -0.255 | 0.822 | 2 | -0.23 (1.58) |
| Time 1 Post –Time 3 Pre | PD | -1.996 | 0.093 | 6 | -1.05 (1.39) |
|  | HC | -- | -- | -- | -- |
| Time 1 Post – Time 3 Post | PD | -1.803 | 0.121 | 6 | 1.14 (2.48) |
|  | HC | -1.973 | 0.299 | 1 | -1.12 (0.80) |
| Time 1 Pre –Time 4 Pre | PD | 1.164 | 0.176 | 4 | 0.84 (.115) |
|  | HC | -- | -- | -- | -- |
| Time 1 Pre –Time 4 Post | PD | 0.936 | 0.402 | 4 | 0.96 (2.28) |
|  | HC | -- | -- | -- | -- |
| Time 1 Post – Time 4 Pre | PD | 1.343 | 0.237 | 5 | 0.68 (1.24) |
|  | HC | -- | -- | -- | -- |
| Time 1 Post – Time 4 Post | PD | 1.197 | 0.285 | 5 | 0.98 (2.00) |
|  | HC | -- | -- | -- | -- |
| Time 3: Pre/Post | PD | 1.565 | 0.149 | 10 | -0.04 (0.91) |
|  | HC | 2.521 | 0.240 | 1 | 1.53 (0.86) |
| Time 3 Pre – Time 4 Pre | PD | 2.102 | 0.069 | 8 | 1.08 (1.55) |
|  | HC |  |  |  |  |
| Time 3 Pre—Time 4 Post | PD | 3.567 | **0.007**** | 8 | 1.90 (1.60) |
|  | HC | -- | -- | -- | -- |
| Time 3 Post – Time 4 Pre | PD | 1.453 | 0.184 | 8 | 0.75 (1.56) |
|  | HC | -- | -- | -- | -- |
| Time 3 Post – Time 4 Post | PD | 3.313 | **0.009**** | 9 | 1.50 (1.44) |
|  | HC | -- | -- | -- | -- |
| Time 4: Pre/Post | PD | 1.009 | 0.337 | 10 | 0.55 (1.80) |
|  | HC | 3.677 | 0.169 | 1 | 1.20 (0.46) |
